# Simplest Model of Nervous System. III. Partial Optimization

**DOI:** 10.1101/2024.06.20.599964

**Authors:** Anton V. Sinitskiy

**Affiliations:** ML LC, Massachusetts, USA

## Abstract

This paper extends our previous work on the simplest model of a nervous system by providing an asymptotic analysis of the evolutionarily optimal solution in this model. Building on the formalism and principles established earlier, we derive an asymptotic solution to the Fokker-Planck-Kolmogorov equation for a given dynamical equation for the state of the neuron. This solution provides the stationary probability distribution for the position of an organism in its environment and the state of its nervous system. Next, exact and asymptotic solutions for the optimal motor and sensory responses to the approach of a predator are derived. These results align with biological expectations and provide a robust mathematical framework for predicting the behavior of simple nervous systems.

## Introduction

In previous work,^1-4^ we presented a novel, highly simplified model of the nervous system inspired by a hypothetical scenario of its origin. Numerical simulations demonstrated that this model successfully captures key properties of the nervous system, and evolutionary optimization changes these properties in the correct direction. For the formulation of the problem, mathematical definitions, discussion of the context, and literature review, we refer the reader to our previous work.^1,2^ Evolutionary optimization in this model can be formalized as the minimization of the functional *I* defined as:

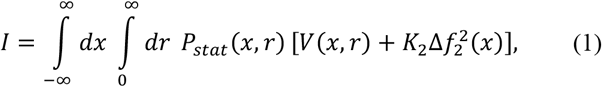

where

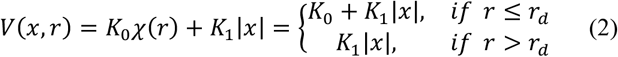

and *P*_*stat*_ (*x, r*) is the stationary probability density, which can be found from the Fokker-Planck-Kolmogorov equation:^5^

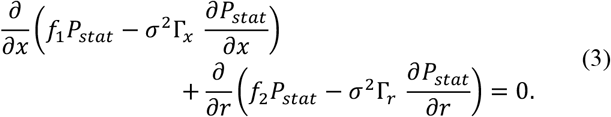

In these equations, *K*_0_, *K*_1_, *K*_2_, *r*_*d*_, *σ*^2^Γ_*x*_, and *σ*^2^Γ_*r*_ are positive constants, *f*_2_(*x, r*) = *f*_*pred*_(*r*) +Δ*f*_2_(*x*), and *f*_*pred*_ (*r*) is a known function. Note that by definition, *I* ≥ 0. For the definitions of all these variables and functions, see our previous work.^1,2^
In general, the functional *I* depends on Δ*f*_2_ [both explicitly, according to equation (1), and implicitly, via *P*_*stat*_ according to equation (3)] and *f*_1_ [only implicitly, via *P*_*stat*_, according to equation (3)]. However, in this work, we perform a partial optimization of *I* as the functional of only Δ*f*_2_, assuming that the function *f*_1_(*x, r*) is known, and has the following form:

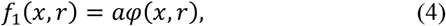

where *a* → +∞ in agreement with the numerical results from our previous work,^1,2^ and φ(*x, r*) is a known function such that φ(*x, r*) = 0 at each *r* has only one solution *x* = *x*_*eq*_ (*r*). We also assume, in agreement with the previous work,^1,2^ that *x*_*eq*_(*r*) possesses the following properties:

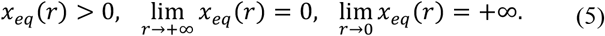

Previously, we considered the case of φ(*x, r*) = −*x* + *C*_*a*_ exp(−λ*r*), where *C*_*a*_ = *C*/*a* stays finite while *a* → +∞.^1,2^ We are interested in the behavior of the solution in the limit of *a* → +∞; while taking this limit, we consider the other functions and constants in this problem, namely *V*(*x, r*), *K*_2_, *σ*^2^, Γ_*x*_, Γ_*r*_, *f*_*pred*_ (*r*), φ(*x, r*), independent of *a*.

### Asymptotic solution to the Fokker-Planck-Kolmogorov equation

For convenience, we will derive the series expansion over *a* not for *P*_*stat*_ (*x, r*), but for the function *u*(*x, r*) defined as:

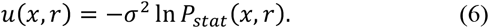

In terms of *u*, equation (3) can be rewritten in the following form:

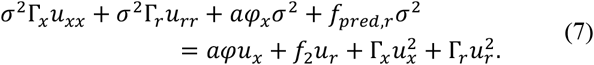

Assume that the scaling of Δ*f*_2_ at *a* → +∞ is *O*(*a*^0^).
If *u*(*x, r*) = *a*^*n*^*S*(*x, r*) + *o*(*a*^*n*^) with *n* > 1, then the leading terms are those of the order *O*(*a*^2*n*^), namely 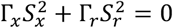, which implies that *S*_*x*_ = *S*_*r*_ = 0; therefore, *S* = *const* and *P*_*stat*_ = *const* · (1 + *o*(*a*^*n*^)), which is not acceptable (contradicts normalization).
Now assume that the leading term is of the order *a*:

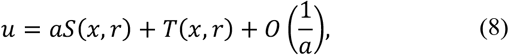

then the leading terms in equation (7) are of the order *O*(*a*^2^), yielding:

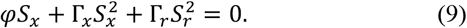

we will search for the solution *S*(*x, r*) in the form of a Taylor series expansion in Δ*x* = *x* − *x*_*eq*_ (*r*) near *x* = *x*_*eq*_ (*r*) for any given *r*:

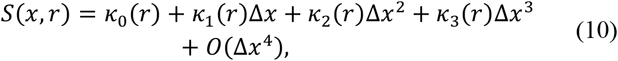

which means that

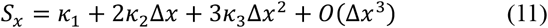

and

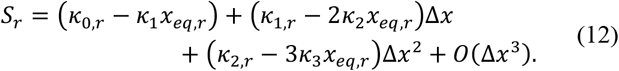

we also expand φ(*x, r*), bearing in mind that at Δ*x* = 0 we have φ(*x, r*) = 0 by the definition of *x*_*eq*_ (*r*):

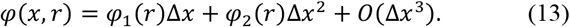

Plugging *S*_*x*_ and *S*_*r*_ into (9) and sorting the terms by the powers of Δ*x*, we get a hierarchy of equations for the coeffictions in the expansions. For the terms of the order Δ*x*^0^ we get:

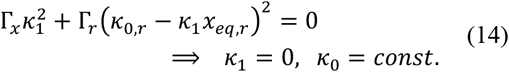

Hence, *S*_*x*_ and *S*_*r*_ simplify to

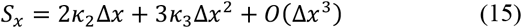

and

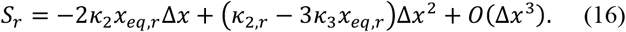

The terms of the order Δ*x*^1^ are now absent. For the terms of the order Δ*x*^2^:

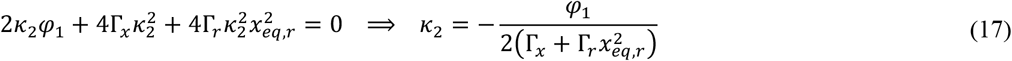

For the terms of the order Δ*x*^3^:

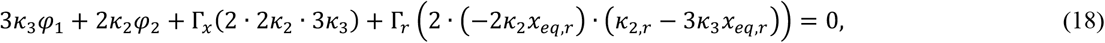

hence

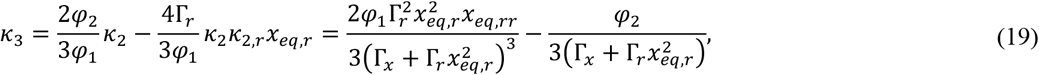

etc. As a result,

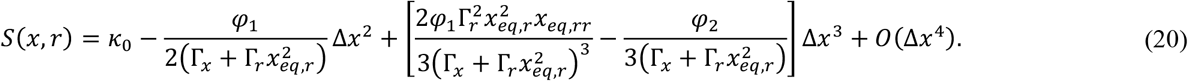

If ∀*r*: φ_1_(*r*) < 0 (recall that in the case for which our numerical simulations were run previously,^1,2^ φ_1_(*r*) = −1, so this condition is always satisfied), then at any *r*, a minimum of *S* as a function of *x* is reached at *x* = *x*_*eq*_ (*r*). Below, we assume that at any *r*, such a minimum is the global minimum over *x*.
As follows from equations (6), (8), and (20),

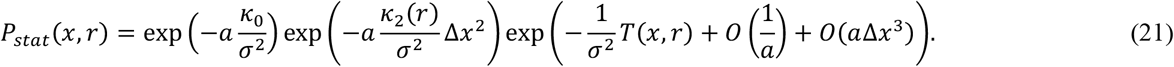

Due to the *O*(Δ*x*^2^) term in *S*, typical values of Δ*x* are of the order 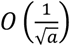, and therefore in the limit of *a* → +∞, *O*(*a*Δ*x*^3^) and higher order terms in *S* vanish in comparison to the *O*(Δ*x*^2^) term for typical values of Δ*x*. Taking into account that a Dirac delta function can be written as a limit

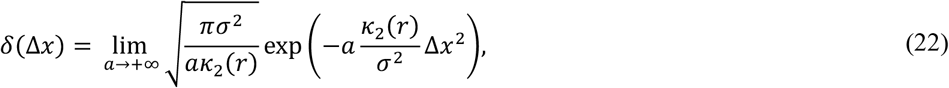

we transform the expression for *P*_*stat*_(*x, r*) to

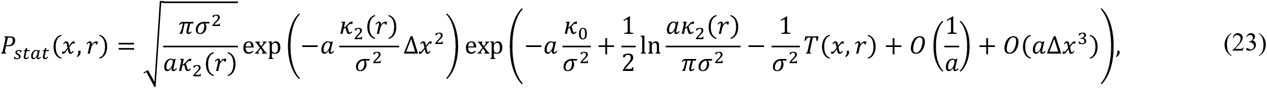

For this expression to converge in the *a* → +∞ limit, we select *κ*_0_ = 0. Also, note that the expansion for *u* in terms of *a*, equation (8), strictly speaking, should also contain a term *O*(ln *a*). This term, however, does not interfere with the differential equation (7) or the hierarchy of equations like (9), (25), etc, because its partial derivatives with respect to *x* or *r* vanish. Hence, in the limit of *a* → +∞, we get

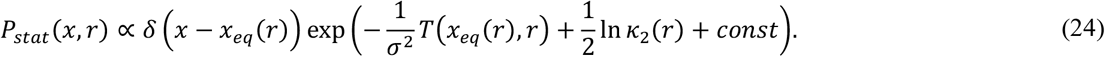

Now return to the expansion of *u* in terms of *a*. The next order leading terms in equation (7) are of the order *O*(*a*), yielding the equation

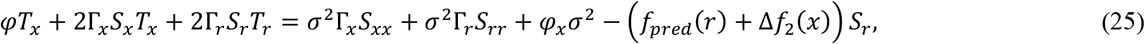

which can be rewritten as

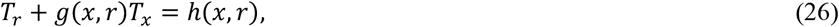

where

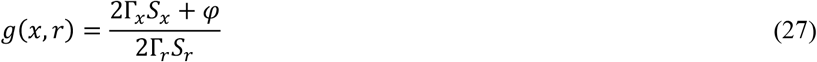

and

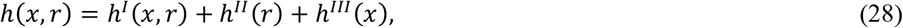

with

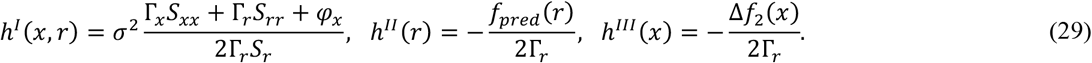

A general solution to this differential equation can be written as

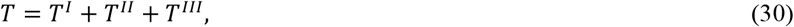

where *T*^*J*^ with *J* = *I, II, III* is a solution to the equation

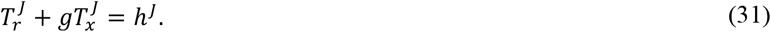

For *T*^*II*^, an exact solution can be written immediately:

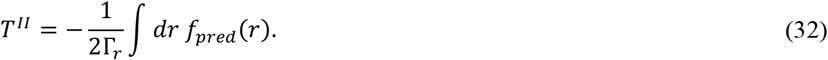

Note that the probability distribution *P*_*stat*_ (*x, r*) in the limit of *a* → +∞, according to equation (24), includes not just *T*, and not the lower-order terms in *a* from the series for *u*, but, moreover, only the value of *T* at Δ*x* = 0. For this reason, let us try to get the expressions for *T*^*I*^ and *T*^*III*^, considering the small Δ*x* series expansions for *g, h*^*I*^ and *h*^*III*^. These series can be obtained from the definitions of these functions, equations (27) and (29), with the use of the expression for *S* given by equation (20):

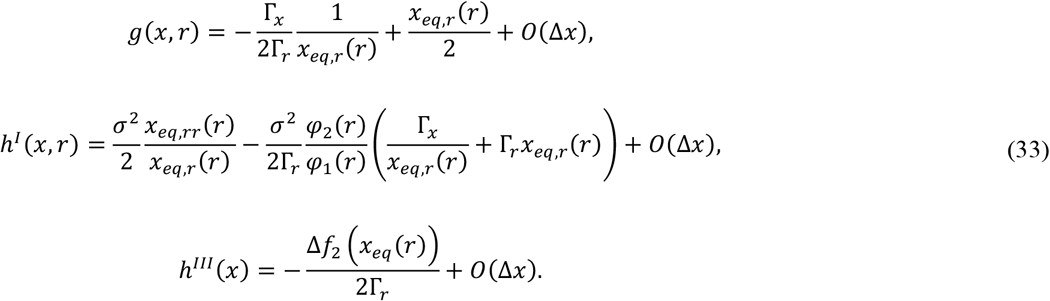

Let us find *T*^*I*^ and *T*^*III*^, defined by equations (31), at Δ*x* = 0, using *g* and *h* given by the leading terms from (33). Assuming that *T*^*I*^ and *T*^*III*^ depend only on *r*, and therefore 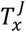 vanishes, we get the following simplest solutions to the differential equation:

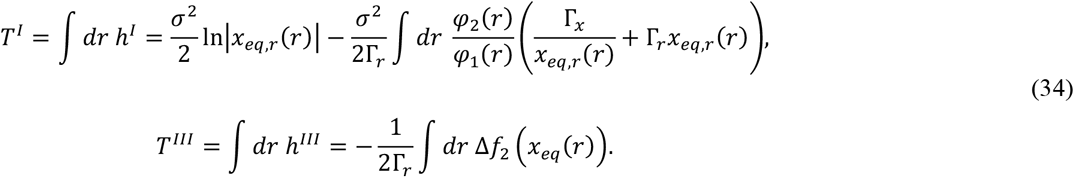

Our numerical simulations^1^ were conducted for *x*_*eq*_ (*r*) = *C*_*a*_ exp(−λ*r*), φ_2_ = 0, φ_1_ = −1, *f*_*pred*_(*r*) = *σ*^2^Γ_*r*_ /*r* − ε*r* and Δ*f*_2_(*x*) = *kx*.
In this case, *T* predicted from equation (34) is

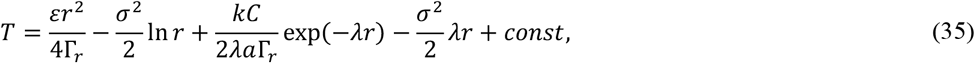

or, with the numerical values of the parameters used for simulations,

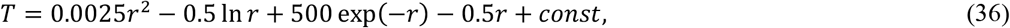

while the distribution *P*_*stat*_ (*x, r*) generated by numerical simulations^1^ corresponds to

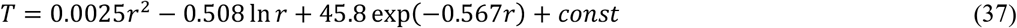

(the linear term in *r* was not included in the fitted functional form because we did not expect it to be there). The similarity of the numerical and analytical (derived neglecting *O*(Δ*x*) terms in *g, h*^*I*^ and *h*^*III*^) expressions is impressive, especially taking into account that the numerical result (37) refers to *a* = 1, not the infinite *a* limit. Another choice of *f*_1_ and *f*_2_, for which exact expressions, without ignoring small in Δ*x* terms, can be written, is provided in Appendix.

### Variational problem for *I* as a one-dimensional integral

Plugging the expression for *P*_*stat*_(*x, r*), equation (24), into the expression for the functional *I* being minimized, equation (1), we can perform integration over *x*, leading to:

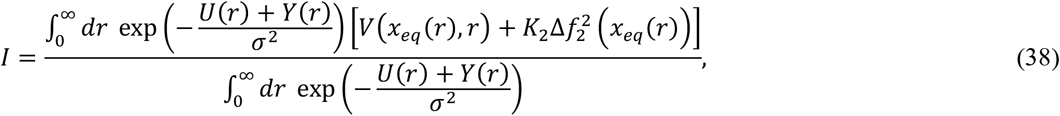

where *U*(*r*) = *T*^*III*^(*x*_*eq*_ (*r*), *r*) is the contribution to *T* coming from Δ*f*_2_(*x*) via *h*^*III*^(*x*), and *Y*(*r*) can be derived from equations (24), (32) and (34):

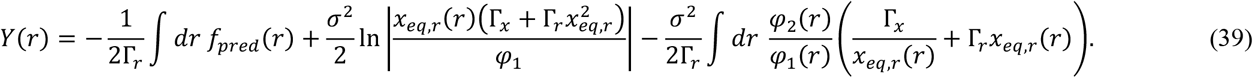

Note that Δ*f*_2_ (*x*_*eq*_ (*r*)) in equation (38) can be expressed from equation (34) in terms of *U*(*r*):

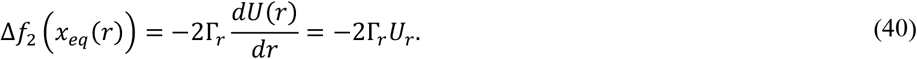

All the other terms in equation (38) are fixed. Therefore, the original problem of minimizing *I*[Δ*f*_2_] given by equation (1) reduces to the problem of minimizing *I* given by equation (38), which can be considered as a functional of *U* only, because both *P*_*stat*_ and Δ*f*_2_ can be found from *U* by equations (24) (exactly) and (40) (probably, in the lowest order in Δ*x*, as assumed in the derivation of equation (34); however, it should be true in general that Δ*f*_2_ =Δ*f*_2_[*U*]).
Therefore, we arrive at the following variational problem: minimize *I*[*U*], given by

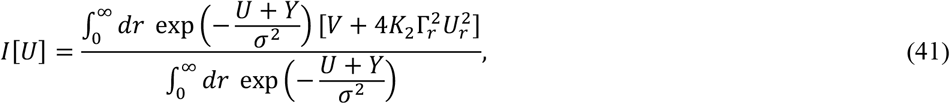

as a functional of *U*. The variation of *I*, given that *U* changes to *U* + δ*U*, equals

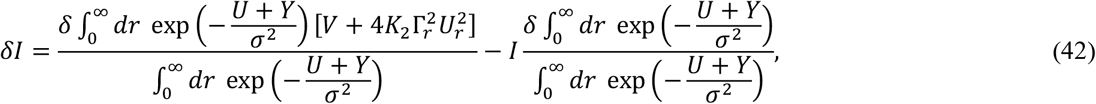

so δ*I* = 0 is reached if

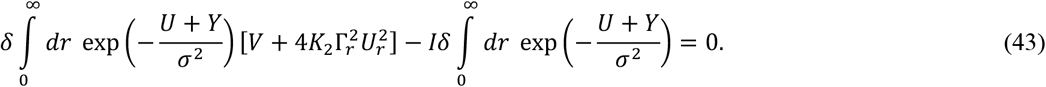

writing the variations explicitly,

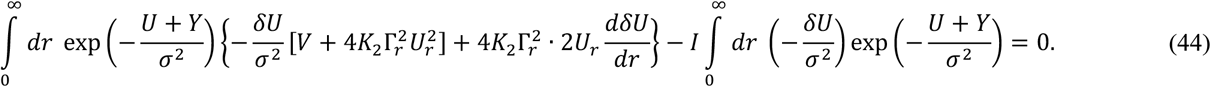

Integration by parts, in the assumption that

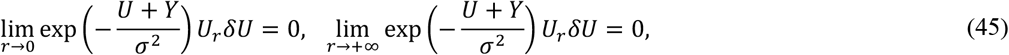

leads to

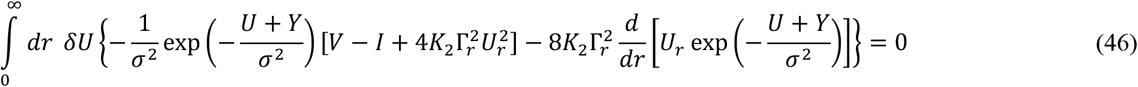

which is satisfied for arbitrary δ*U* only if

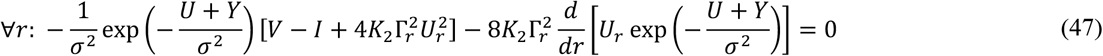

that is

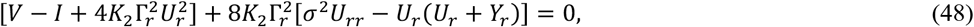

leading to

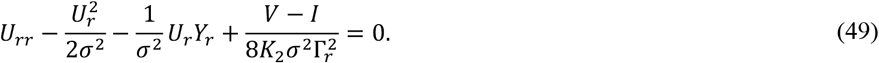

Making a substitution

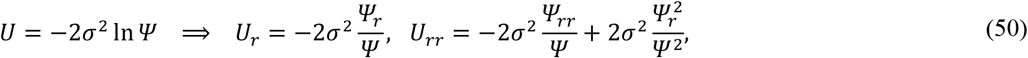

we get the differential equation

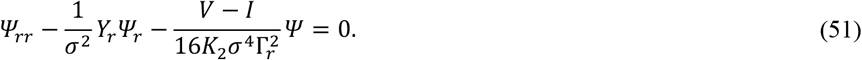

Using the expression for *Y* given by equation (39), we get

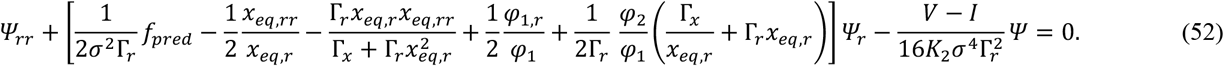

In the limit of *r* → +∞, and therefore *x*_*eq*_(*r*) → 0^+^, and assuming that, as in numerical simulations,

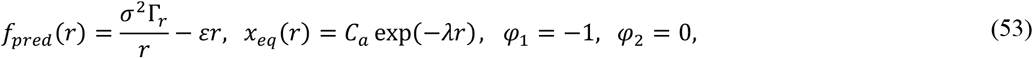

we get the following asymptotic behavior of the coefficients in the differential equation:

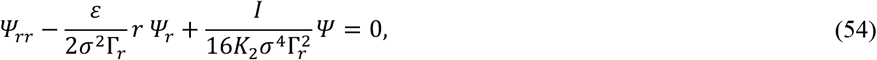

and therefore the following asymptotic behavior of the solution at *r* → +∞

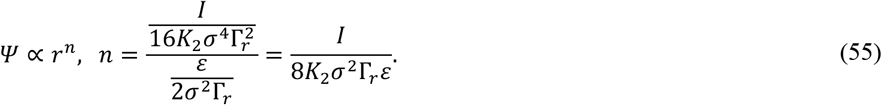

Then we can get the following asymptotic for *U*:

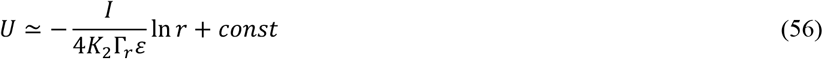

and from equation (40),

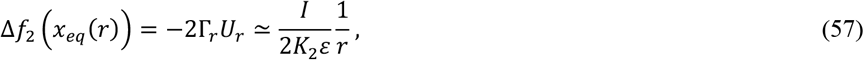

therefore,

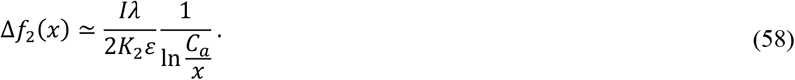

Note that

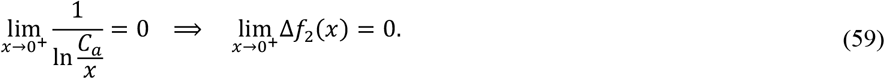

The asymptotic behavior of the differential equation (52) at *r* → 0^+^, with the same assumptions (53), is:

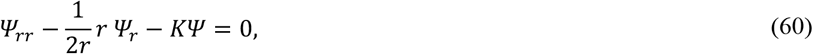

where:

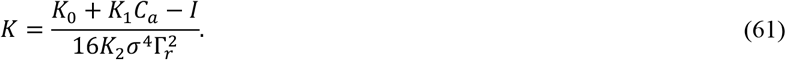

Therefore, the asymptotic behavior of the solution at *r* → 0^+^ is

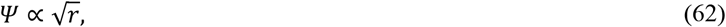

which corresponds to

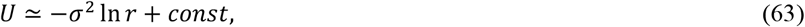

and from equation (40),

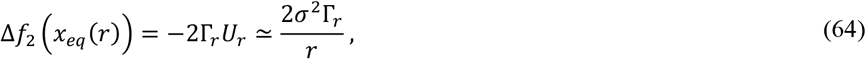

therefore,

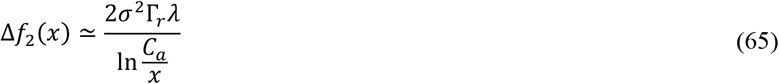

and Δ*f*_2_ → +∞ in the considered limit. (The other independent solution of equation (60) leads to Ψ ∝ exp(*Kr*^2^/3) and 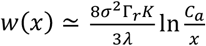, which tends to zero in the considered limit of *r* → 0^+^, which is not compatible with the interpretation of the problem.)
From the viewpoint of the biophysical interpretation, asymptotes (57) and (64) are totally reasonable. When the nearest predator is far (*r* → +∞), the motor reaction is weak (Δ*f*_2_ → 0; note also that this happens not only because of *r* → +∞, but also the prefactor is proportional to *I*, so for an optimized solution it is small). When the predator approaches (*r* → 0^+^), sharp motor response is triggered (Δ*f*_2_ → +∞; this time, the prefactor is independent of *I*). Interestingly, both asymptotes (57) and (64) have the same functional form (inversely proportional to *r*), but with different prefactors. As we will see below, the value of *I*, and therefore the prefactor in the asymptote for *x* → 0^+^, can be made arbitrarily small. On the other hand, the prefactor in the asymptote for large *x* is a finite constant 2*σ*^2^Γ_*r*_. Therefore, we can expect that the exact Δ*f*_2_(*x*) undergoes a ‘stepwise’ increase at some intermediate *x*, switching from one hyperbolic curve to another one, lying higher.
For the purpose of further derivations, it is not necessary to use the exact solutions of equation (52). An approximate solution, demonstrating roughly the same qualitative behavior of Δ*f*_2_(*x*), will suffice for the purpose of minimizing *I*.

### Value of the functional *I* with a qualitatively reasonable approximation for Δ*f*_2_

Here we will find the functional *I* for the following specific choice of Δ*f*_2_(*x*):

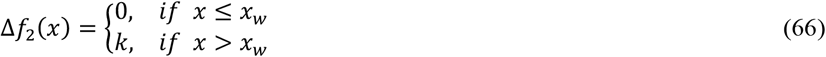

where *k* is a large positive constant, and *x*_*w*_ is some positive constant (the threshold value of the potential, such that the motor reaction turns on only when the nervous system excites above this threshold level). This choice corresponds to a qualitative understanding of what the optimal Δ*f*_2_(*x*) look like, as discussed at the end of the previous section. Then

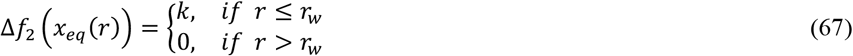

where *r*_*w*_ is such that *x*_*eq*_ (*r*_*w*_) = *x*_*w*_. Hence, choosing the integration constant in equation (34) in such a way that *U*(*r*_*w*_) = 0, we get

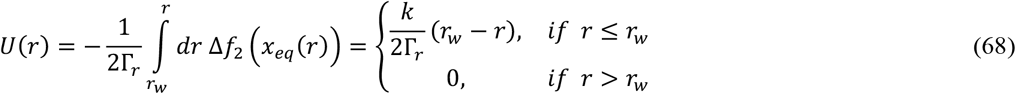

Next, we split the integral of exp(− (*Y* + *U*)/*σ*^2^) in (41) into two parts,

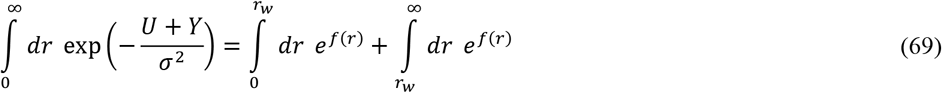

where, in the case given by equation (53),

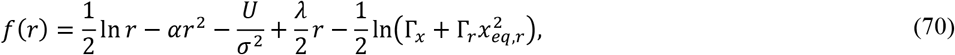

where we introduced a short-hand notation:

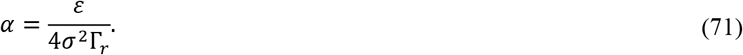

First, consider the case of *r* > *r*_*w*_. Then *U* = 0 and

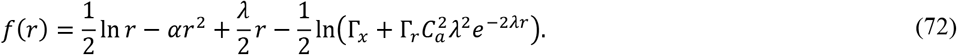

The maximum value of *f*(*r*) is achieved at

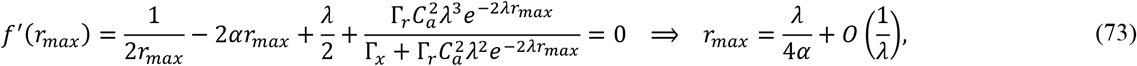

where

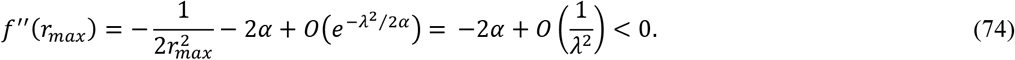

confirming that this is a maximum. Thus, at large λ, *r*_*max*_ maximizing *f*(*r*) grows as λ/4α. Similarly, the value of the argument that maximizes 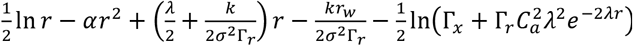, which is the functional form of *f*(*r*) at *r* ≤ *r*_w_, is also on the order of λ/4α. Thefore, at sufficiently large λ, this value of *r* will exceed *r*_*w*_, and the only maximum of *f*(*r*) is achieved at the value of the argument given by equation (73), which also satisfies

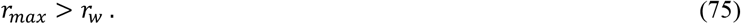

Therefore, the second integral in equation (69) can be approximated as

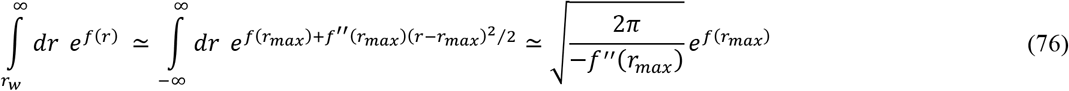

which in the large λ limit can be estimated, with the use of equations (70), (73) and (74), as

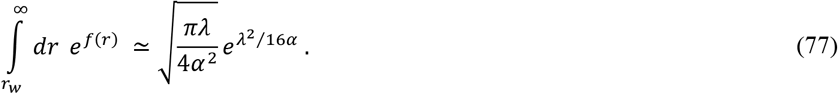

To estimate the first integral in equation (69), note that at *r* ≤ *r*_*w*_, and therefore at *r* < λ/4α,

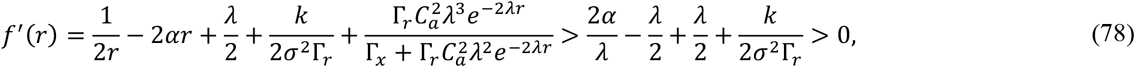

so the maximal value of *f*(*r*) within 0 ≤ *r* ≤ *r*_*w*_ is reached at *r* = *r*_*w*_. Then, expanding *f*(*r*) near *r* = *r*_*w*_, the integral of *e*^*f*(*r*)^ can be estimated as:

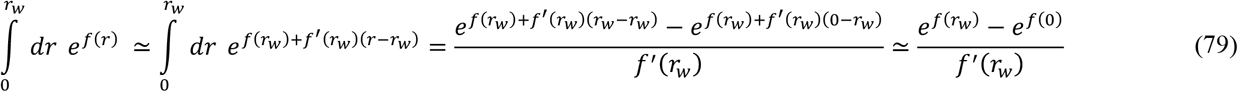

which in the large λ limit can be estimated, with the use of equations (70), (73) and (74), as

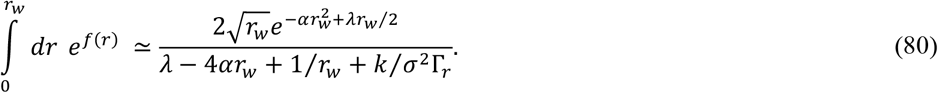

Note that the ratio of the two integrals *R* defined as

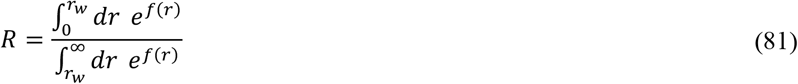

in the large λ limit tends to zero:

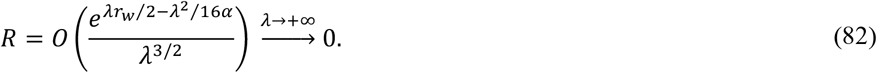

This result makes sense from the viewpoint of the biophysical interpretation of the problem, because greater sensitivity of the neural system to the approaching predator (λ → +∞) means that the most probable distance to the nearest predator *r*_*max*_ will increase, and the probability that the nearest predator approaches close to the organism (*r* ≤ *r*_*w*_) will get smaller and smaller (*R* → 0). Then the components of the functional *I* can be estimated as follows:

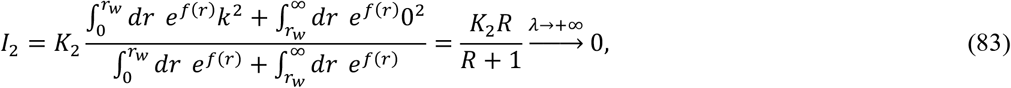

next,

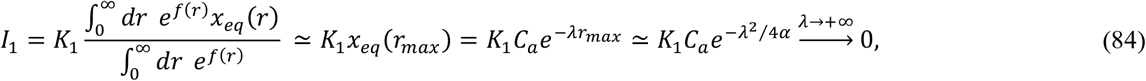

and finally, assuming that *r*_*d*_ ≤ *r*_*w*_ (the distance to the nearest predator at which the escape behavior is triggered is not smaller than the distance at which the organism can be eaten),

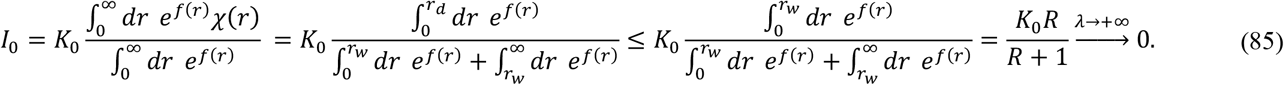

Hence, even though we chose Δ*f*_2_(*x*), equation (66), not exactly in the form of an exact solution to equation (52), nevertheless the whole functional *I* can be done arbitrarily close to zero, which is its lowest possible value, because from the definition *I* ≥ 0, by choosing λ to be large enough:

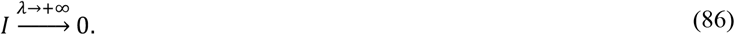

Thereby, we have found a particular solution to the initial problem of minimizing *I*. Specifically, the value *I* can be made arbitrarily close to 0 in the following case:

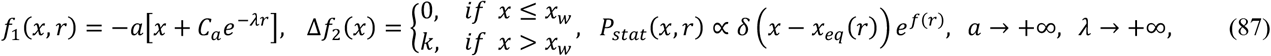

where *f*(*r*) is defined by (70), *x*_*w*_ is a positive constant not exceeding 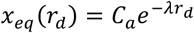, and all other parameters (including *C*_*a*_ and *k*) are arbitrary positive constants.

### Optimal solution for the function *x*_*eq*_

In previous sections, we assumed a specific functional form of *x*_*eq*_, namely *x*_*eq*_ (*r*) = *C*_*a*_ exp(−λ*r*). In this section, we lift this assumption, while keeping in force the other assumptions from equation (53), as well as the choice of Δ*f*_2_(*r*) given by equation (67), and explore what functional form of *x*_*eq*_(·) would be optimal.
In principle, we could consider the functional *I*, equation (38), as a functional of *x*_*eq*_ (·). This time, *U* is fixed due to the choice of Δ*f*_2_(*r*) (Δ*f*_2_(*x*) will depend on *x*_*eq*_(·), because Δ*f*_2_(*x*) =Δ*f*_2_ (*r*_*eq*_ (*x*)), but this dependence does not affect the minimization of *I*[*x*_*eq*_]). The effect of *x*_*eq*_ on *I* is via *Y*(*r*), which depends on *x*_*eq*,*r*_ according to equation (39), and via *V*(*x*_*eq*_(*r*), *r*), which depends on *x*_*eq*_. Our attempt to derive *x*_*eq*_ explicitly via the Euler–Lagrange equation led to a second-order differential equation with a complicated nonlinear dependence on *x*_*eq*_ and its first derivative, an equation that we could not solve.
However, we notice that a quantitative consideration can be performed by analogy with the analysis in the previous section. In the case of an arbitrary *x*_*eq*_, the function *f*(*r*) assumes the following form:

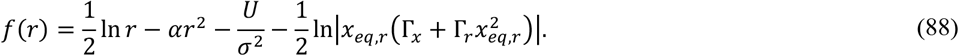

At *r* > *r*_*w*_, *U* = 0 and therefore

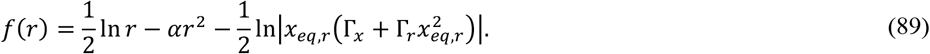

we expand the log term in equation (88) in Taylor series in the vicinity of some distance *r*_*ref*_ that will be specified later:

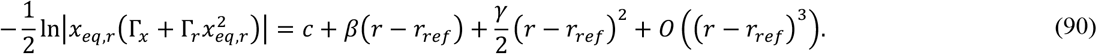

Plugging this into the expression for *f*(*r*), we get

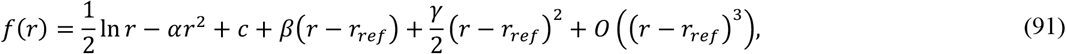

which has the maximum at *r*_*max*_ given by

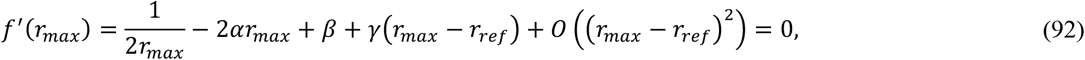

provided that the second derivative at this point

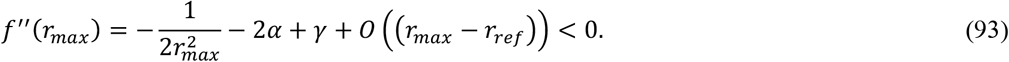

Now we identify *r*_*ref*_ with this *r*_*max*_ to get

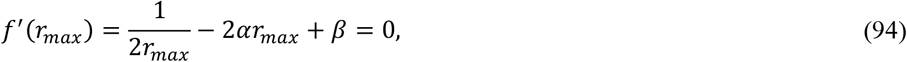

which, taking into account that *r*_*max*_ > 0, leads to the following solution for *r*_*max*_ :

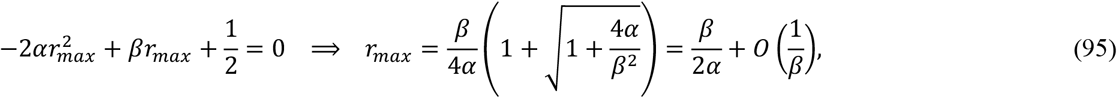

and the following restriction on *γ*:

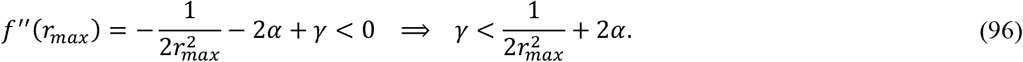

Similar to equation (76), we can compute

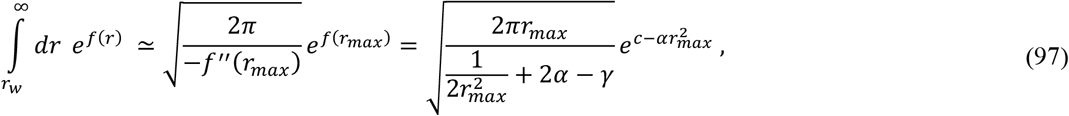

which for large *β* can be estimated as

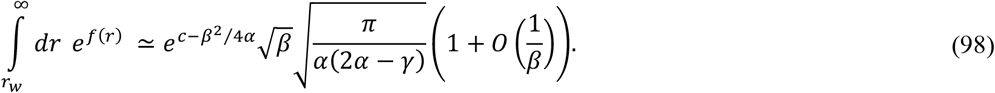

Now consider the case of *r* ≤ *r*_*w*_. To compute the other integral, by analogy with equation (79), we expand the log term in equation (88) in Taylor series in the vicinity of *r*_*w*_; the coefficients in the series this time will be different:

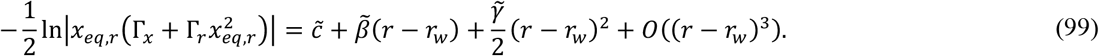

Then the full function *f*(*r*) at *r* ≤ *r*_*w*_ is

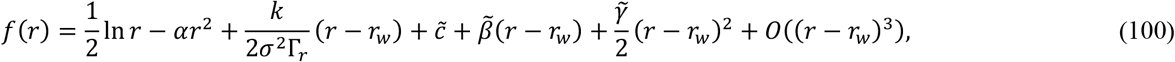

and its first derivative at *r*_*w*_

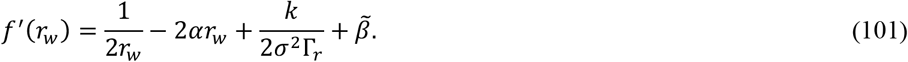

Because our goal, as in the previous section, is to minimize the ratio *R*, it is reasonable to assume that

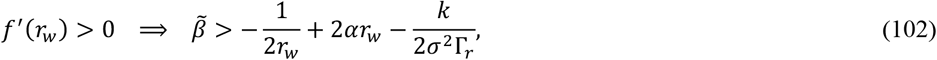

otherwise the integral 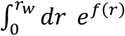 will get higher contributions from *e*^*f*(*r*)^ at some *r* < *r*_w_. Then, similar to equation (79),

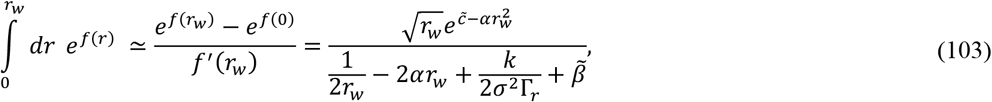

and the ratio *R* can be expressed as

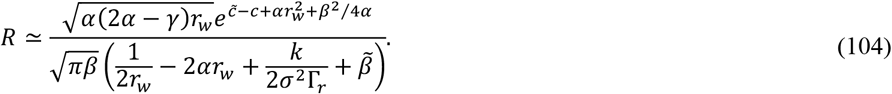

As in the previous section, the contributions *I*_0_ and *I*_2_ to the functional *I* being minimized are bound above by the following expression:

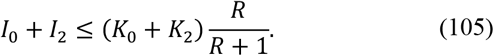

Let us first minimize *I*_0_ + *I*_2_, and then we will check that the resulting conditions also ensure *I*_1_ → 0, and therefore, *I* → 0. Minimization of *I*_0_ + *I*_2_ can be achieved with *R* → 0. Notice that a sufficient condition for *R* → 0 is 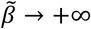, while all other parameters 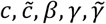 may stay finite. This is not the only way to get *R* → 0. Other scenarios include 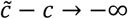 with finite 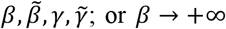 provided that 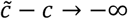 such that 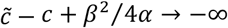. Also, increasing *γ* closer to its maximal allowed value as given by equation (96) decreases *R*, though not to zero; however, increasing *γ* also deteriorates the accuracy of approximation (97). A faster approach of *R* → 0 can be achieved with a combination of these trends, for example,

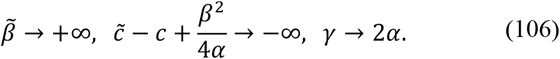

Now, consider what kind of *x*_*eq*_(·) could ensure this solution. For the limit of 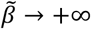, equation (99) implies that in the leading order

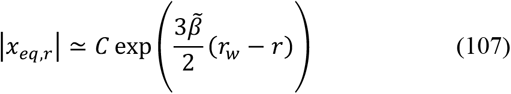

grows rapidly (in the limit, infinitely rapidly) as *r* → *r*_*w*_. This can correspond to two variants for |*x*_*eq*_ (*r*)|: it should either grow or decay rapidly (in the limit, infinitely rapidly) as *r* → *r*_*w*_. Out of these two variants, based on the biophysical interpretation, we choose the variant of rapid decrease of |*x*_*eq*_ (*r*)| in this region.
To minimize the remaining contribution *I*_1_ to the funcitonal *I*, we also assume that |*x*_*eq*_(*r*)| at *r* < *r*_*w*_ stays finite (which implies that the series expansion (100) is not applicable at *r* noticeably smaller than *r*_*w*_):

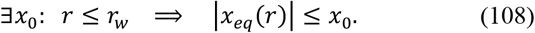

Then we can get the following restriction:

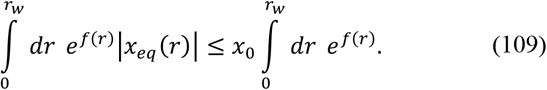

This restriction could possibly be made less strict, and |*x*_*eq*_ (*r*)| slowly increasing to infinity as *r* → 0, such that *e*^*f*(*r*)^|*x*_*eq*_ (*r*)| decreases to zero fast enough, would also work.
Now, consider the integral of *e*^*f*(*r*)^|*x*_*eq*_ (*r*)| from *r*_*w*_ to +∞. Due to the multiplier |*x*_*eq*_ (*r*)|, the position of the maximum of the integrand shifts from *r*_*max*_ given by equation (95) to a somewhat different value *r*_*max*1_. Then, the value of the integral can be evaluated in the leading order as

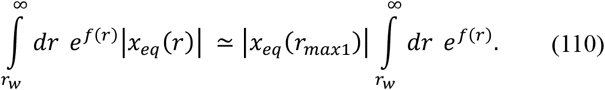

Then, the upper bound on the contribution *I*_1_ to the functional *I* being minimized can be found as follows:

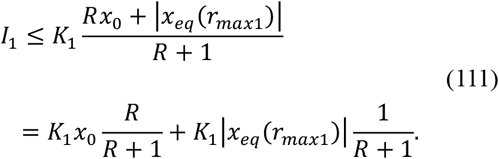

The first term on the right hand side of equation (111), similar to the upper bound on *I*_0_ + *I*_2_ in equation (105), tends to zero because *R* → 0. As for the second term, in order for it to vanish, *x*_*eq*_ (*r*_*max*1_) has to vanish. This can be achieved if *x*_*eq*_ (·) quickly decays while *r* increases from *r*_*w*_ to *r*_*max*1_. In the previous section, this decay was exponential, but, as the analysis in this section shows, any other function quickly decaying to zero would also work.
Thus, we have reproduced the final conclusion of the previous section, that the optimal *x*_*eq*_ (*r*) stays finite at *r* < *r*_*w*_ and quickly (in the limit of λ → +∞, infinitely quickly) decreases with the growth in *r*. However, the analysis in this section reveals more details on what general conditions are imposed on *x*_*eq*_(*r*), and how they affect achieving the desired result of *I* → 0.
The fast (in the limit, infinitely fast) growth of *x*_*eq*_ (*r*) with decreasing *r* near *r* = *r*_*w*_ – from the viewpoint of the biophysical interpretation, infinitely high sensitivity of the nervous system at the border of the danger – is critical for a small (in the limit, infinitely small) probability for the organism to find itself in the dangerous zone *r* ≤ *r*_*d*_, and therefore, nullification of *I*_0_, *I*_2_, and the contribution to *I*_1_ that appears from being in this zone. Additionally, this effect can be enhanced by a rapid decay of *x*_*eq*_ (*r*) not only at *r* = *r*_*w*_, but also in the range of *r* between *r*_*w*_ and *r*_*max*_ or *r*_*max*1_, as expressed by the condition of 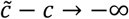, and a faster-than-exponential growth of *x*_*eq*_ (*r*) with decreasing *r* near *r* = *r*_*w*_, as suggested by the condition *γ* → 2α (implying *γ* > 0). However, as *r* → 0 at *r* < *r*_*w*_, *x*_*eq*_(*r*) should not infinitely increase (at least, fast enough) to ensure that the vanishing probability of being in the dangerous zone is not overcome by an even faster increase in *K*_1_|*x*_*eq*_ (*r*)|. As for the region of *r* > *r*_*w*_, *x*_*eq*_ (*r*) should decay fast in this region. This ensures that the main contribution to *I*_1_, which is the cost of the work of the nervous system under typical circumstances in the absence of the danger, vanishes because of the vanishing excitation of the nervous system in this safe zone.

## Discussion

The paper presents an analytical (specifically, asymptotic) analysis of the previously proposed simplest model of the nervous system.^1,2^ Minimization of a functional that represents evolutionary fitness of an organism with the nervous system is performed in several steps. First, an asymptotic solution to the Fokker-Planck-Kolmogorov equation has been derived, providing the stationary probability distribution for the system (the organism in its environment). This probability is shown to be sharply peaked in one-dimensional slices of this two-dimensional distribution function. Based on this result, the original problem of minimizing a two-dimensional integral is reduced to a one-dimensional problem, and its exact solution and asymptotic behavior at large and small distances to the predator is explored. Additionally, the optimal solution for the function *x*_*eq*_ (·) describing the sensory role of the neuron is presented. The analysis provides insights into the optimal response of the simplest nervous system to stimuli. It shows that a sharp response is triggered when a predator is close, while the response weakens as the predator moves away. This behavior aligns with biological expectations, where the motor reaction should be stronger when the organism is in immediate danger.
The main value of this work is in the proposed robust mathematical framework for understanding the probabilistic behavior of a simple nervous system. The derived solutions and conditions offer predictive power for how neurobiological systems might behave. Additionally, the methodology can be used to optimize artificial systems mimicking biological responses, where similar decision-making processes take place.
A few open questions remain. In particular, it is not clear why if we take Δ*f*_2_ = 0 (that is, we turn off the motor reaction of the organism), then the potential *Y*, according to equation (39), still depends on *x*_*eq*_ (*r*). In this case, the organism only feels an approaching predator, but does not escape it. It makes sense that in this case the full probability distribution *P*_*stat*_ (*x, r*) depends on *x*_*eq*_ (*r*), because the nervous system feels the predator approach, hence the distribution over *x* changes. However, the integral of the distribution over *x*, that is 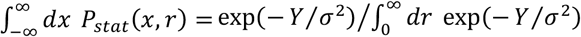, which describes only the distance to the predator *r*, should not depend on these internal sensations, because they do not result in any motor reaction. Hence, according to the biophysical interpretation of the problem, *Y*(*x*_*eq*_ (*r*), *r*) for Δ*f*_2_ = 0 should not depend on *x*_*eq*_ (*r*), in contradiction to equation (39). Similarly, it is not clear why the choice of *x*_*eq*_ (*r*) near the maximum of *f* affects 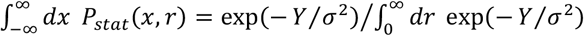, as quation (89) suggests, even if Δ*f*_2_(*r*) is chosen in the form of a stepwise function, equation (67). The motor reaction at these distances does not turn on, hence, a slightly higher or lower excitation of the neuron should not change the distance to the predator *r*. Another open question is whether the derivation of equation (34), and therefore expressions for *U* and *Y*, is justified with the approximation of *g, h*^*I*^ И *h*^*III*^ by their values at Δ*x* = 0, as given by equations (33), and what happens to the term in *T* that corresponds to the solution to the homogenous equation *T*_*r*_ + *g*(*x, r*)*T*_*x*_ = 0 (see also Appendix for a particular case that may help to clarify this issue). These questions need further investigation.
In conclusion, we have presented an analytical solution to the partial problem of evolutionary optimization in the previously proposed simple model of the nervous system. This solution is obtained explicitly through an asymptotic expansion. Unlike previous studies based on optimization of a few numerical parameters, the derivation in this paper yields general functional forms of the optimal neuronal and motor reactions. This analysis may pave the way for understanding complex nervous systems in light of the evolutionary approach, without depending on historic frozen accidents and particular material implementations of nervous systems.

## Acknowledgements

I would like to thank Prof. S.V. Shaposhnikov for valuable discussions. I am solely responsible for any mistakes in this work.

## Appendix

Here we consider the solution to the Fokker-Planck-Kolmogorov equation for a specific choice of functions *f*_1_ and *f*_2_, qualitatively reproducing the key properties of these functions revealed in the analysis of the main text:

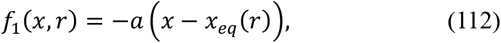

where

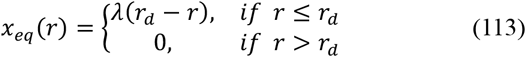

and

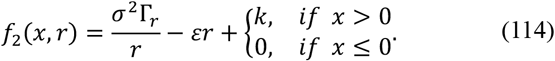

Here λ is a new positive constant, while all other constants are the same as earlier. This choice of *f*_1_ and *f*_2_ ensures that the sensory system recognizes an approach of a predator only at close distances, and the motor reaction turns on only if the neuron is excited, while in the absence of excitation it does not work and therefore saves energy.
At *r* > *r*_*d*_, *f*_1_ = −*ax* and *f*_2_ = *σ*^2^Γ_*r*_ /*r* − ε*r*, so an exact solution can be written:

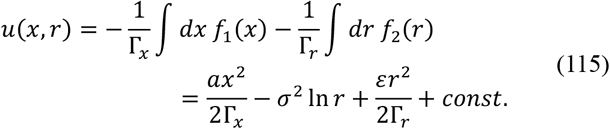

At *r* ≤ *r*_*d*_, we find the solution in a form given by equation (8). Then, equation (9) assumes the following form:

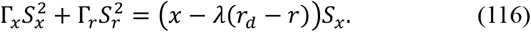

In the series for *S* in terms of Δ*x*, the third-order and higher-order terms vanish, and the exact solution for *S* can be written as follows:

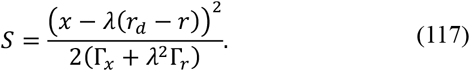

From equation (25), the following exact differential equation for *T* can be obtained:

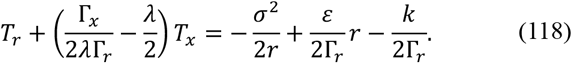

A general solution for *T* can be written as a sum of a particular solution to this equation and a solution to the corresponding homogeneous differential equation. One particular solution can be easily found if we assume that *T* depends only on *r*, but not *x*. Then

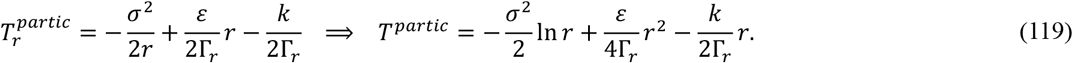

A solution to the homogenous equation 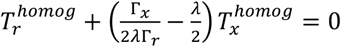 can be found by the method of characteristics, yielding

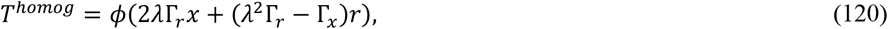

where ϕ(·) is an arbitrary differentiable function. Therefore, the general expression for *T* is

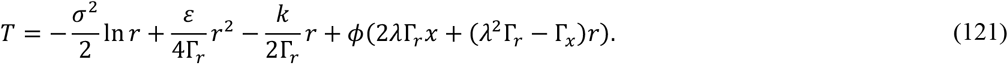

It is not quite clear to us yet how to choose this function ϕ(·). One variant would be, as in the main text, to set ϕ = 0, which presumably ensured agreement with our previous numerical simulations in that particular case. Another variant might be to get ϕ(·) from the condition of continuity of *u*(*x, r*) at *r* = *r*_*d*_, the border at which the two solutions, equation (115) and (121) [in the second case, taking into account (117)], should coincide:

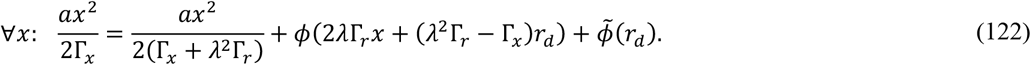

Here, the last term 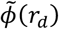 is a collection of all the terms in equations (115) and (121) that do not depend on *x*. By making a substitution:

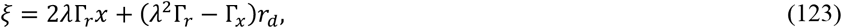

we can rewrite equation (122) as:

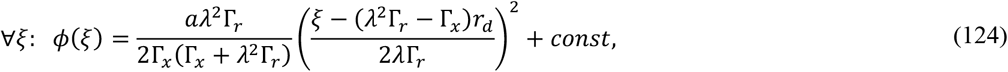

where the constant does not depend on *r* or *x*, and therefore, equation (121) as

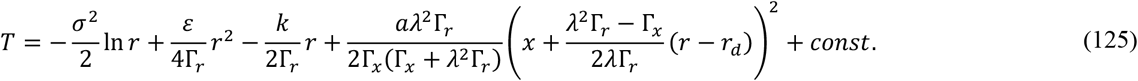

This expression, however, is problematic in that it contains the parameter *a* that was used for the series expansion of *u*, while the term *T* by construction, equation (8), should not contain *a*. Perhaps, equation (115), to which we matched equation (121), is not a (unique) solution for *u*(*x, r*) at *r* > *r*_*d*_. These questions might be resolved by numerical simulations with *f*_1_ and *f*_2_ given by equations (112)-(114), by analogy with our previous work.

## Notes

### Competing Interest Statement

The authors have declared no competing interest.

